# Long-term non-invasive drug treatments for adult zebrafish

**DOI:** 10.1101/2021.12.21.473492

**Authors:** Yuting Lu, E. Elizabeth Patton

**Affiliations:** MRC Human Genetics Unit and CRUK Edinburgh Centre, Institute of Genetics and Cancer, University of Edinburgh, Western General Hospital Campus, EH4 2XU, Edinburgh, UK

**Keywords:** adult zebrafish, drug pellets, non-invasive, long-term treatment, melanoma, vemurafenib, melanoma regression, drug resistance models

## Abstract

Zebrafish embryos are widely used for drug-discovery however administering drugs to adult zebrafish is limited by current protocols that can cause stress. Here, we develop a drug formulation and administration method for adult zebrafish by producing food-based drug pellets which are consumed voluntarily. We apply this to zebrafish with BRAF-mutant melanoma, a model that has significantly advanced our understanding of melanoma progression, but not of drug resistance due to the limitations of current treatment methods. Short-term, precise, and daily dosing with drug-pellets made with the BRAF^V600E^ inhibitor, vemurafenib, led to tumour regression. On-target drug efficacy was determined by phospho-ERK staining. Continued drug treatment led to the emergence, for the first time in zebrafish, of acquired drug resistance and melanoma relapse, modelling the responses seen in melanoma patients. This method presents a controlled, non-invasive approach that permits long-term drug studies, and can be widely applied to any adult zebrafish model.

**SUMMARY STATEMENT:** This drug-pellet approach is a precise dosing method to administer short or long-term drug treatments to adult zebrafish, that is stress-free for the fish and requires minimal animal handling. We use this method to develop new drug-resistant models of melanoma in zebrafish, opening new doors for modelling and screening drug treatments in adult zebrafish.

## INTRODUCTION

Over the past two decades zebrafish have emerged as an important model for drug discovery, directly leading to drugs that have entered clinical trials or for compassionate use (Patton et al., 2021b). These include therapies that promote hematopoietic stem cell renewal (North et al., 2007), prevent antibiotic-induced ototoxicity (Kitcher et al., 2019), and those that treat childhood epilepsy (Baraban, 2021), cancer (Mandelbaum et al., 2018; White et al., 2011; Yan et al., 2019), lymphatic anomaly (Li et al., 2019), arteriovenous malformation (Al-Olabi et al., 2018), and fibrodysplasia ossificans progressive (Yu et al., 2008), among others. This success is, in part, because zebrafish are vertebrates and share over 80% of disease genes with humans (Howe et al., 2013), as well as shared drug metabolism, physiology and pharmacology (MacRae and Peterson, 2015). Thus, zebrafish pre-clinical disease models are important platforms for drug discovery and repurposing, even at times leading to new treatment strategies for patients directly from zebrafish models.

With some exceptions, most zebrafish drug discovery, gene-drug screens and compound-phenotype evaluation studies are performed using zebrafish embryos. However, embryos and larval stages do not fully recapitulate adult disease states and lack a complete immune system. Drug screening and discovery in adult zebrafish – and modelling the impact of long-term drug treatment – has been limited by methods of drug administration, which can be complex, harmful, and imprecise. Current methods for experimental drug discovery in adult zebrafish involve adding the drug to the fish water which can irritate exposed mucosal surfaces (*e.g*. eyes, gills), is not appropriate for water-insoluble compound administration, and involves administrating large quantities of drug to fish water with unknown final drug concentrations absorbed by the fish. Alternatively, drugs can be administered by oral gavage or through injection (retro-orbital or intraperitoneal), which are more precise for dosing, but invasive and sometimes fatal, and require repeated anesthesia (Dang et al., 2016; Kinkel et al., 2010; Pugach et al., 2009). While these methods are generally acceptable for short-term drug treatments, they are problematic for longer term drug protocols because they can lead to accumulative distress and injury to the animal. Adding antibiotics to jelly like food has been used to manage zebrafish colony health (Chang et al., 2017), however, these methods were not designed to administer precise drug treatments to individual fish in the experimental and pre-clinical disease model context, and are therefore not appropriate to investigate dose-based drug responses. Thus, non-invasive and precise drug delivery protocols that permit longitudinal experimental drug treatments for adult zebrafish are not well developed.

Our laboratory uses zebrafish to model the progression of melanoma, the deadliest form of skin cancer (Travnickova and Patton, 2021). Zebrafish melanoma models have provided key insights into the origins, progression and new drug targets for melanoma (Baggiolini et al., 2021; Ceol et al., 2011; Kaufman et al., 2016; Patton et al., 2005; Travnickova et al., 2019; Venkatesan et al., 2018; White et al., 2011). However, a significant gap in the field has been to generate zebrafish melanoma models that recapitulate the development of acquired drug resistance as seen in patients (Patton et al., 2021a). Indeed, acquired resistance is one of the major challenges limiting the progression-free survival time of melanoma patients on therapies that specifically target BRAF^V600E^ and MEK signalling (Chapman et al., 2011; Larkin et al., 2014; Long et al., 2015; Luke et al., 2017; Ribas et al., 2014; Robert et al., 2015; Sosman et al., 2012). This gap is unfilled in zebrafish melanoma models due to the lack of sustainable long-term drug administration methods for adult fish, despite evidence for the strong potential of zebrafish models recapitulating many human melanoma plasticity states, including residual disease (Travnickova and Patton, 2021).

Here, we present a novel drug formulation and administration method for adult zebrafish which enables the delivery of precise drug concentrations directly via food pellets. As a proof-of-principle, we fed zebrafish with BRAF^V600E^ melanomas pellets containing vemurafenib and showed that they caused immediate and on-target efficacy in reducing melanoma growth. Long-term treatments (>2 months / daily treatment) led to drug resistance and melanoma progression, enabling zebrafish models of melanoma drug resistance for the first time. Our drug pellet method opens new doors to model the effects of drug dose, administration, and long-term treatments in adult zebrafish that is non-invasive and limits animal handling, and is widely applicable across a wide range of zebrafish disease models.

## RESULTS

### Formulation of drug-pellets for adult zebrafish

We wanted to generate drug pellets that would deliver consistent drug doses to treat melanoma while also being quickly and freely consumed by the zebrafish. To begin, we prepared vemurafenib pellets using a drug-pellet mould that we designed and 3D-printed, so that each drug pellet would be 2 mm in diameter (comparable to the size of a zebrafish egg, and which are easily consumed by adults) **(Figure 1A)**. We suspended dry fish food in water, and mixed this with agar-agar and gelatine powder to create a food paste to generate the base of the food pellet. To prepare the pellets, 10 cm^3^ dry fish food was added to water up to 50 ml in a conical centrifuge tube, completed with 1 g of food-grade agar-agar plus 2 g of gelatine powder, and shaken well to generate a red-coloured mix **(Figure 1B)**. This recipe was optimised to achieve a balance between congelation for laboratory handling and a palatable texture for the fish. The mixture was then transferred into a 100 ml borosilicate container and microwaved for ~1 min to reach boiling, turning the mixture into a smoothie-like texture **(Figure 1B)**. Just before the mixture congealed, we added the desired drug (*i.e*. vermurafenib) or DMSO as a control **(Figure 1A)**. We prepared our vemurafenib pellets to each deliver 100 mg/kg vemurafenib (as shown to induce tumour regression when administered by oral gavage)(Dang et al., 2016). Briefly, 10 mg of vemurafenib powder was resuspended in 300 μl DMSO and then mixed well with 700 μl agar-fish food mixture while warm in a 1.5 ml microcentrifuge tube, generating a pink-coloured paste **(Figure 1C)**. Each well of the 3D-printed pressing mould is 5 μl, and therefore each pellet contained 0.05 mg vemurafenib. Given that the average weight of each fish is 0.5 g, one vemurafenib pellet per day will deliver the dose of 100 mg/kg.

**Figure 1.**
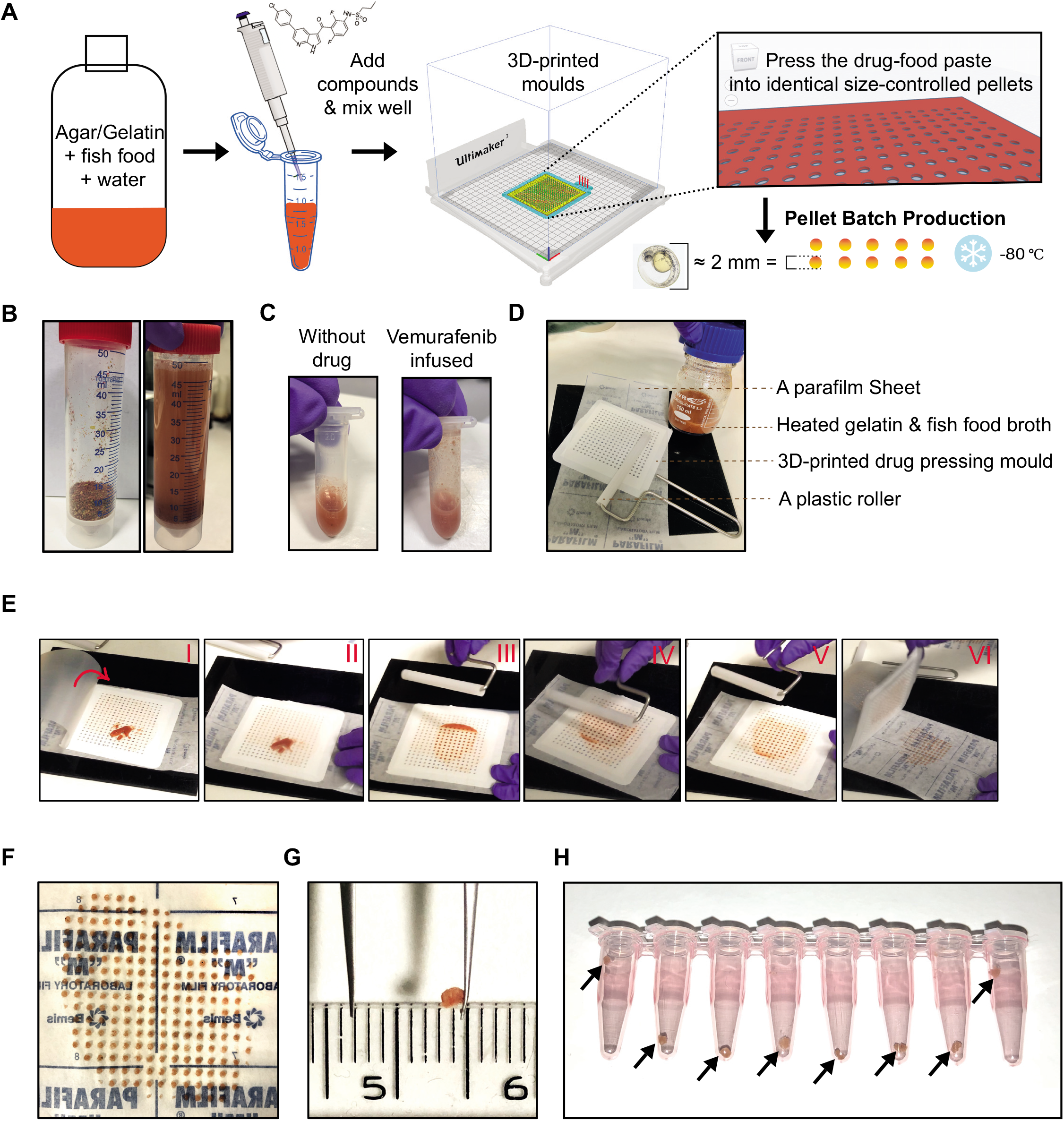
Technical setup and drug pellet preparation. **A.** Schematic overview of the drug-pellets formulation and manufacturing pipeline. **B.** Dry fish food mix and the food-agar mixture resuspended in water. **C.** The red-colour food-agar paste becomes pink once supplemented with vemurafenib dissolved in DMSO. **D.** The tools used for pressing drug-pellets in the mould. **E.** Sequential series of photos (I-VI) showing the process of drug-pellet pressing. The parafilm sheet is peeled from the backing paper, and the mould is placed on the backing paper. The drug-paste is applied on to the mould (I), and the parafilm sheet is gently lowered to cover the paste and mould (II). Next, using the roller, the drug-paste is evenly applied into the holes of the mould (III-V). The parafilm sheet is lifted, followed by carefully removing the mould, and the drug pellets adhere to the backing paper (VI). **F.** A freshly prepared batch of drug-pellets. Surface tension retains the pellets on the parafilm backing paper. **G.** A drug-pellet recovered from −80°C storage, maintaining the flat-cylinder shape. The ruler shown in the picture is scaled in cm/mm. **H.** Drug-pellets aliquoted into daily doses per fish in PCR tubes, ready for −80°C storage, with arrows highlighting the drug-pellets inside the tubes.

A sheet of parafilm large enough to cover the 3D-printed drug pressing mould and an appropriately sized plastic roller were prepared as tools to press drug pellets **(Figure 1D)**. Once the food-drug paste began to congeal, the paste was transferred onto the drug-pressing mould, covered with the parafilm sheet (to form a “sandwich” with the mould between the parafilm sheet and the backing paper), and then pressed into the mould with the plastic roller **(Figure 1E)**. The parafilm sheet was lifted, and the mould carefully removed to reveal the drug pellets now formed and adhering to the backing paper (surface tension will trap most of the drug pellets neatly on the backing paper) **(Figure 1E-G)**. This approach enables batches of >100 cylindrical tablet-like drug pellets to be conveniently made during a single preparation. DMSO control pellets were prepared in a similar way. In this experiment, drug pellets were prepared once every week and stored at − 80°C as individual pellets in PCR tubes for daily dose aliquots (**Figure 1H**). The entire process of pressing drug pellets is shown in **Video 1**.

### Administration of drug-pellets to adult zebrafish by free feeding

We then fed the drug-pellets to individual adult zebrafish with melanoma that were singly-housed in 1 L tanks. As shown in **Video 2**, zebrafish actively sought for and consumed the drug pellet voluntarily. In general, zebrafish consumed their drug pellets immediately, however, any pellets that were ignored for more than 15 minutes were removed and replaced by fresh pellets to ensure consistent dosing in drug administration. Using a pipette to gently deliver the drug pellets encouraged zebrafish to spot and consume the pellets. We observed no distress or toxicity in the fish from either the drug-pellet formulation or the administration.

### Vemurafenib drug-pellets reduce zebrafish BRAF^V600E^ melanoma burden

To test the drug efficacy, we provided DMSO or vemurafenib drug pellets to adult zebrafish with BRAF^V600E^ *p53* mutant melanomas (Patton et al., 2005) and compared the tumour responses between the two groups. To track the tumour size over time, we imaged and analysed the brightfield images of each tumour from these individuals every week **(Figure 2A)**. DMSO treated zebrafish showed continuous lesion expansion and tumour growth **(Figure 2B, 2C)**, while 100 mg/kg vemurafenib treated fish showed tumour regression over 3 weeks **(Figure 2D, 2E)**, with the average tumour size reduced by 60% in 2 weeks, and 70% in 3 weeks, compared to the pretreatment state **(Figure 2E).** This result is highly comparable to the observation from the previous 100 mg/kg oral-gavage regime (Dang et al., 2016), from which they reported 2-week daily treatment reduced the melanoma size by 70% in average.

**Figure 2.**
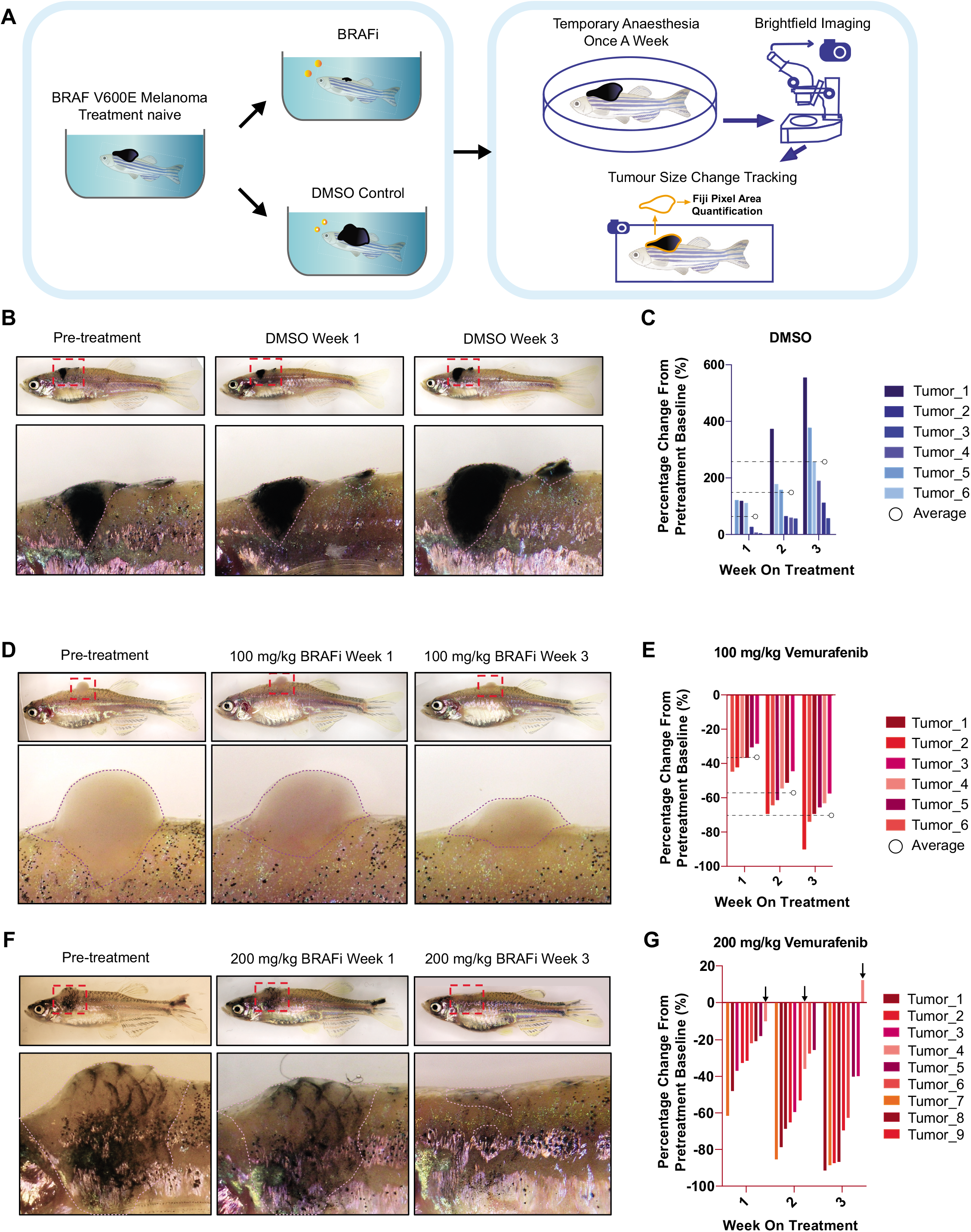
Short-term assessment of vemurafenib pellets on BRAF^V600E^ zebrafish melanoma. **A.** Schematic overview of drug-pellet free feeding administration, tumour response tracking, and evaluation for each fish treated with vemurafenib or DMSO pellets. **B.** Representative images of BRAF^V600E^ zebrafish melanoma progressing on treatment of DMSO pellets. Zoomed regions are indicated by red dashed boxes. Dotted line outlines the melanoma. **C.** Quantification of melanoma size change each week (by percentage) on DMSO pellets, comparing to the lesion imaged on the day pre-treatment. Fish receiving DMSO pellets (N=4; tumour counts n=6). Average of tumour size changes indicated by dashed lines. **D.** Representative images of BRAF^V600E^ zebrafish melanoma regressing while treated with 100 mg/kg vemurafenib pellets. Dotted line outlines the melanoma. **E.** Quantification of melanoma size change each week on 100 mg/kg pellets by percentage, compared to pre-treatment. Fish receiving 100 mg/kg vemurafenib pellets N=3. Lesion counts n=6. Average of lesion size changes indicated by dashed lines. **F.** Representative images of BRAF^V600E^ zebrafish melanoma regressing on treatment of 200 mg/kg vemurafenib pellets. **G.** Quantification of melanoma size change each week on 200 mg/kg pellets by percentage, comparing to the lesion imaged on the day pre-treatment. Fish receiving 200 mg/kg vemurafenib pellets N=4. Lesion counts n=9. One lesion (Tumour 4) with regrowth observed by week 3 on 200 mg/kg vemurafenib pellets is indicated by arrows.

Next, we increased the dose of vemurafenib by feeding the fish 2 drug-pellets per day, and observed a higher proportion of lesions had improved regression (4 out 9 reduced by > 80% in 3 weeks, compared to 1 out 6 in fish treated with 1 pellet/day). Thus, drug-pellets are highly effective at treating zebrafish melanoma, well tolerated, and can be used in dose escalation studies.

### Long-term drug treatments lead to drug resistance

Patients with BRAF mutant melanoma and receiving targeted therapy will often show a dramatic reduction in melanoma burden, followed by recurrent melanoma growth from residual disease (or persister cells)(Marine et al., 2020; Shen et al., 2020b). We and others have shown that persister cell states are heterogeneous and consist of cell states that pre-exist in the primary tumour and emerge *de novo* (Rambow et al., 2018; Travnickova et al., 2019). We have studied these states in zebrafish by conditional expression of the master melanocyte transcription factor MITF, while others have used mouse xenograft studies to follow human melanoma cells following BRAF inhibitors (Rambow et al., 2018; Shen et al., 2020a; Travnickova et al., 2019). However, there are no zebrafish models of BRAF inhibitor resistance over time because of the limitations of long-term drug delivery methods to adults.

We noticed that one tumour developed resistance and regrowth at the end of week 3 on treatment at 200 mg/kg in the short-term treatment protocol (**Figure 2G**; Tumour 4). We reasoned that drug-pellets could be used to investigate the effect of long-term vemurafenib treatment on BRAF^V600E^ zebrafish melanoma, and generate models of drug-resistance. Similar patterns of tumour responses were observed in mouse xenograft models which showed that human melanoma tumours regressed following daily MAPK-inhibitor treatment, and then entered a stable or residual disease stage around 18 days, followed by recurrent growth as drug-resistant tumours around 50 days (Rambow et al., 2018; Wang et al., 2018).

We first treated zebrafish on the course of 100 mg/kg vemurafenib daily pellets, and followed the tumour response over 5 weeks **(Figure 3A, B)**. In our experiment, we noticed that by 4 weeks at 100 mg/kg, the melanomas began to regrow, so we increased the dose to 200 mg/kg at week 5 **(Figure 3B)**. We found that melanomas were initially responsive to the increased vemurafenib concentration, but that they again began to regrow by 8 weeks **(Figure 3B)**. Next, we treated a cohort of zebrafish with melanomas with 200 mg/kg vemurafenib for four weeks, and found that the melanomas responded rapidly to the treatment, with drug resistance and progressive disease observed as early as 4 weeks **(Figure 3C, 3D)**. These studies indicate that our drug-pellet delivery methods can model long-term treatment response adaptation, from the initial melanoma regression, through to stable disease, and finally recurrent and drug resistant disease.

**Figure 3.**
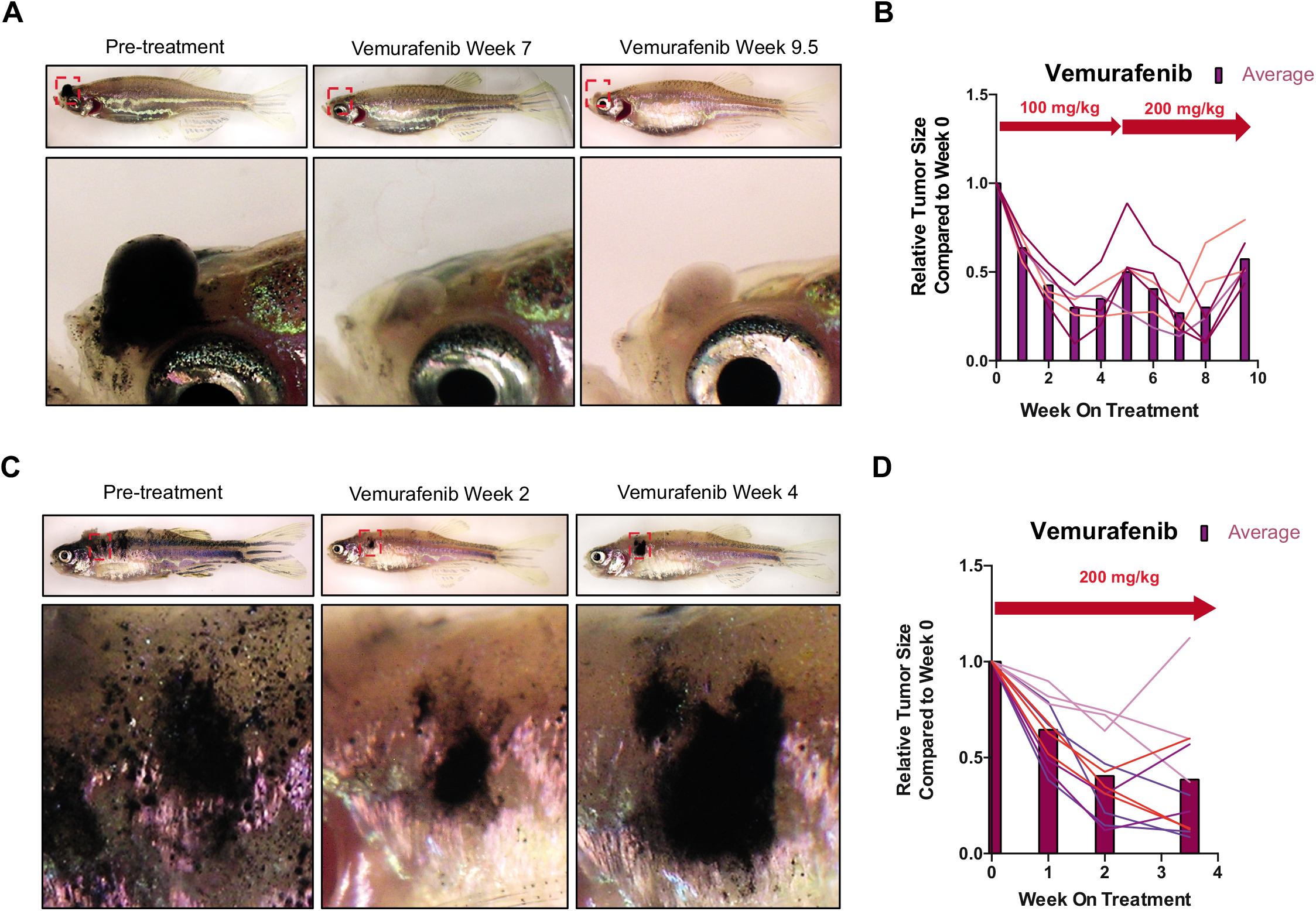
Long-term vemurafenib drug pellet treatment causes acquired drug resistance in zebrafish melanoma. **A.** Representative images of BRAF^V600E^ zebrafish melanoma before treatment, regressed melanoma, and progressive disease for the animals shown in **B**. **B.** Quantification of melanoma size change each week on 100 mg/kg vemurafenib pellets or 200 mg/kg after dose escalation. Fish receiving vemurafenib pellets N=4. Lesion counts n=6. Each coloured line represents one lesion with the size change tracked over the entire treatment course. Lesions from the same fish share the same colour. The average of all lesions across samples are indicated by column bars. **C.** Representative images of BRAF^V600E^ zebrafish melanoma before treatment, during melanoma regression, and evidence of recurrent disease while on consistent treatment of 200 mg/kg vemurafenib. **D.** Quantification of melanoma size change each week on 200 mg/kg vemurafenib pellets. Fish receiving vemurafenib pellets N=4. Lesion counts n=12. Each coloured line represents one lesion with the size change tracked over the entire treatment course. Lesions from the same fish share the same colour. The average of all lesions across samples are indicated by column bars.

### Validation of drug-pellet efficacy in adult zebrafish cancer

To assess the on-target efficacy of vemurafenib to inhibit BRAF^V600E^ activity in zebrafish melanoma, we performed immunofluorescence staining on melanoma sections to assess the MAPK pathway activity using phospho-Erk1/2, a downstream target of activated BRAF signalling. Melanoma samples collected from the early-responding stage of vemurafenib pellet treatment (weeks 2 and 3) had significantly reduced levels of phospho-Erk1/2 compared to DMSO treated control samples **(Figure 4A, 4B)**, indicating that vemurafenib pellets have sufficient bio-availability to target BRAF^V600E^ in melanoma and to lead to melanoma regression.

**Figure 4.**
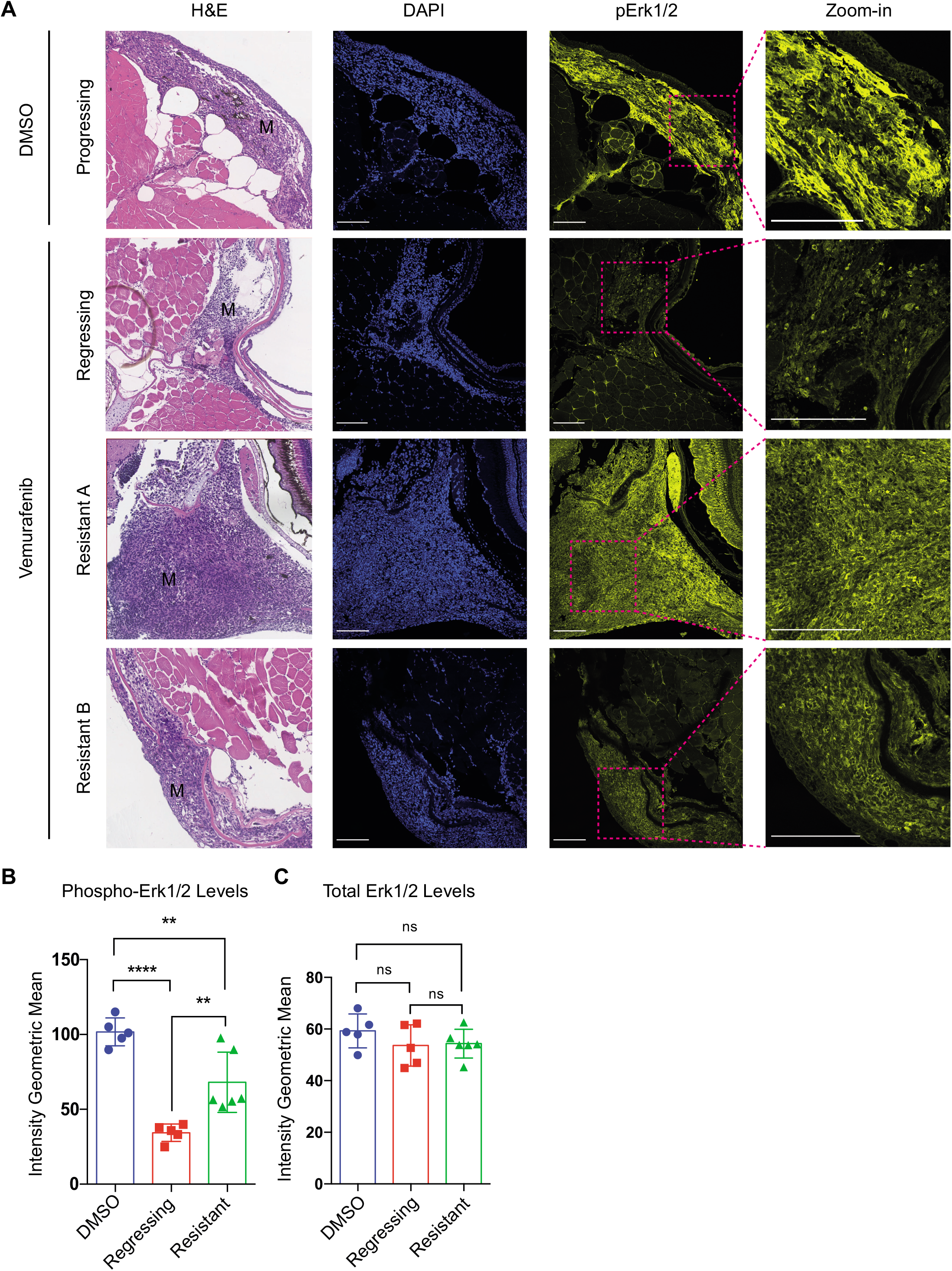
On-target efficacy of vemurafenib drug pellet treatment. **A.** Representative images of H&E and immunofluorescence staining of BRAF^V600E^ zebrafish melanoma samples treated with DMSO or Vemurafenib drug pellets. Phospho-Erk1/2 staining in melanoma cells (M) is clearly visible in zoomed regions. Regressing melanomas have reduced phospho-Erk1/2 staining, and the response is varied in Vemurafenib resistant disease. Scale bar = 100 μm. DMSO treated melanoma sample (week 3, DMSO treatment); melanoma regression sample (week 3, 200 mg/kg Vemurafenib treatment); Melanoma resistant tumour A and B (week 10; 5-week 100 mg/kg, followed by 5-weeks 200 mg/kg Vemurafenib treatment). **B, C**. Quantification of immunofluorescence staining intensity of phospho-Erk1/2 and total Erk1/2 from BRAF^V600E^ zebrafish melanoma samples treated with DMSO, regressing on Vemurafenib drug pellets, and resistant to Vemurafenib. The DMSO treated samples were collected post 2- or 3-week treatment (N=4 fish, n=5 lesions). The regressing samples were collected week 3, 200 mg/kg treatment (N=4 fish, n=5 lesions). The resistant samples were collected week 10, 5-weeks 200 mg/kg Vemurafenib treatment escalation course following the initial 5-week 100 mg/kg Vemurafenib treatment (N=3 fish, n=6 lesions). (Multiple t-test with Sidak-Bonferroni correction. ns, not significant. **p value <0.01; ****p < 0.0001)

Next, we analysed tumours that showed melanoma recurrence on vemurafenib pellets, and found that although the responses were varied, on average, tumours increased levels of phospho-Erk1/2, consistent with that seen in patients (Manzano et al., 2016; Proietti et al., 2020) **(Figure 4A, 4B)**. Total Erk1/2 levels measured by immunofluorescence staining showed no significant changes across all samples **(Figure 4C)**. These results validate the on-target efficacy of the drug compound delivered by our pellet feeding method.

## DISCUSSION

Zebrafish are a powerful model system for drug discovery, but while drug treatments for embryos and larvae can be easily administered through the water, drug discovery in adult zebrafish is limited by a lack of efficient, non-invasive and long-term permissive drug administration methods. Here, we provide a new method to generate drug pellets that can be easily fed to adult zebrafish to administer a controlled and precise drug dose in a non-invasive process. We apply this method to our zebrafish BRAF mutant melanoma models (Patton et al., 2005) and demonstrate that the melanomas respond to vemurafenib drug-pellet therapy. We validate the on-target efficacy of the drug by showing a reduction in total phospho-Erk1/2 in the melanoma following treatment. Longterm studies (>2 months) demonstrate that upon drug treatment, zebrafish melanomas undergo regression followed by recurrent disease, as seen in patients (Marine et al., 2020; Shen et al., 2020b; Travnickova and Patton, 2021). For the first time, this method enables us to model longterm melanoma drug treatment and resistance stages in zebrafish genetic melanoma models, an immunocompetent model system.

We found no toxicity or side-effects from the drug-pellet method, indicating that this method supports the tenets of the 3Rs (Replacement, Reduction and Refinement) in Animal Research (https://nc3rs.org.uk/). Specifically, by reducing the animal handling and total exposure required for drug administration, our method is a Refinement for drug delivery because it minimises zebrafish stress and improves welfare.

In conclusion, we provide a drug-pellet method to administer precise doses of drugs to adult zebrafish in a non-invasive, free-feeding based procedure. The drug-pellets can be individually frozen so that experiments can be controlled for batch effects, are suitable for drugs with low solubility in water (such as vemurafenib, which is hydrophobic), and provides a platform for drug combinations and screens. For a broader spectrum of application, pellet size and number can be adapted easily through modifying the drug-press 3D printing mould and modifying the quantities of the food paste recipe. While our experiments here focus on cancer studies in zebrafish, we expect this method will be applicable to a wide range of zebrafish disease models, and opens new doors for drug discovery in the complex biological context of adult zebrafish.

## RESOURCES TABLE

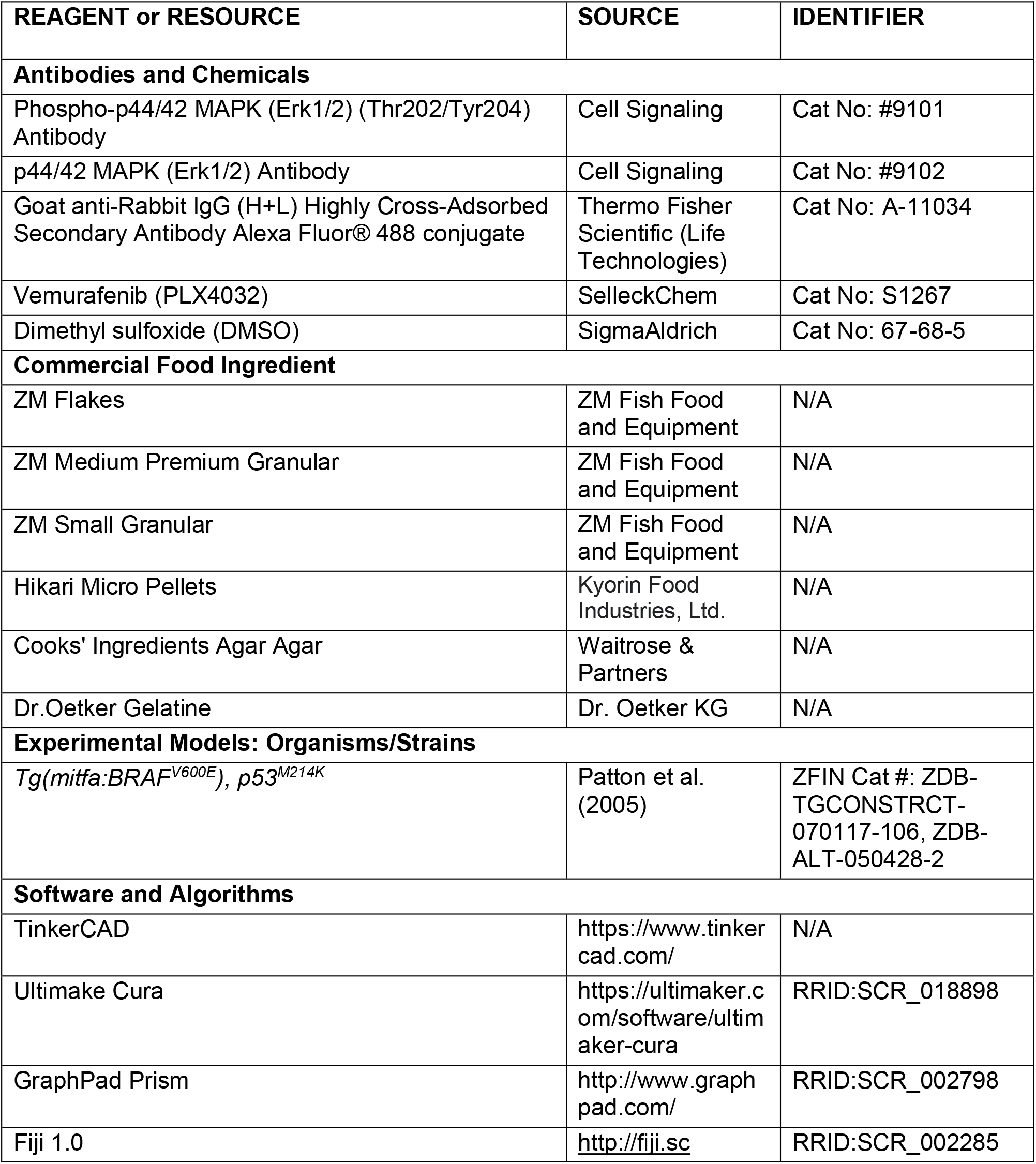

## METHODS

### Zebrafish maintenance and husbandry

Zebrafish were maintained in accordance with UK Home Office regulations, UK Animals (Scientific Procedures) Act 1986, under project license P8F7F7E52. All experiments were approved by the Home Office and AWERB (University of Edinburgh Ethics Committee).

### Zebrafish melanoma models

Zebrafish were genotyped using DNA extracted from fin clipped tissue by PCR to establish the mutant allele status *tp53^M214K^* or *mitfa-BRAF^V600E^* as described in our previous publications (Travnickova et al., 2019). The emergence of melanoma is usually observed in individuals aged 3- to 6-month-old. Individuals used for DMSO versus vemurafenib drug pellets treatment in this experiment were siblings and were aged 5- to 6-month-old when entering the treatment scheme.

### Drug pellets ingredients

The recipe of our routine dry fish food mix consists of ZM flakes, ZM Medium Premium Granular, ZM Small Granular and Hikari MicroPellets mixed at a ratio of 2:3:2:5. Food-grade agar-agar or gelatine powder were purchased from local grocery stores. Vemurafenib (SelleckChem, CAS#918504-65-1) powder were resuspended in DMSO before mixing with fish-food paste as described.

### 3D-printing

The 3D-modelling and design of the drug-pressing mould was carried out on Tinkercad **(Supplementary File 1)** followed by slicing set-up using Ultimaker Cura and 3D-printed via Ultimaker 3 with AA 0.25 generic PLA.

### Imaging of adult zebrafish and tumour size measurement

Fish were briefly anesthetised (Tricaine in PBS 1:10,000 concentration) once every week for imaging purposes to follow the tumour burden changes during the experiment. Each fish was anaesthetised for no longer than 10 min per session and fully recovered in fresh system water. Brightfield images were taken for each fish positioned on both sides. Images of fish lesions were captured at the same magnification scale at the same microscope every week. The size of each lesion was quantified by using the manual field selection in Fiji on each tumour image, then compared to the matching pre-treatment lesion to calculate the relative percentage change. Lesions that could be observed from both sides of the fish were measured by combining the area number averaged from both sides.

### Zebrafish histology and IHC quantification

Zebrafish melanoma samples were collected, fixed, and processed as described in our earlier publications (Lister et al., 2014). MAPK activity was assessed using phospho-p44/42 MAPK (ERK1/2) (Thr202/Tyr204) primary antibody (1:200, rabbit, Cell Signaling Technologies #9101), total p44/42 MAPK (ERK1/2) primary antibody (1:200, rabbit, Cell Signaling Technologies #9102), and Alexa-fluor 488 secondary antibody (1:1000, goat-anti-rabbit IgG, Life Technologies #A-11034). Nuclei were stained with DAPI dye (1:1000, Life Technologies #62248).

## ACKNOWLEDGEMENTS

We are grateful to Cameron Wyatt and the IGC Zebrafish Facility for zebrafish management and husbandry, the IGC Imaging Facility for supporting the imaging experiments, and Helen Caldwell and Elaine McLay for histology. We appreciate the constructive inputs from Cameron Wyatt and Jana Travnickova during the development of this project. We are grateful for uCreate Studio team of the University of Edinburgh for providing the equipment and materials for the 3D-printing self-service. EEP is funded by MRC HGU Program (MC_UU_00007/9), the European Research Council (ZF-MEL-CHEMBIO-648489), and Melanoma Research Alliance (687306).

## AUTHOR CONTRIBUTIONS

Conceptualization: YL; Methodology: YL; Validation: YL; Formal analysis: YL; Investigation: YL; Resources: YL, EEP; Writing original draft: YL, EEP; Writing review and editing: YL, EEP; Visualization: YL, EEP; Supervision: EEP; Funding acquisition: EEP.

## VIDEO LEGENDS

**Video 1. Preparation of drug pellets**.

Drug supplemented food-agar mixture paste was cooled in a petri-dish, and once partially congealed, the jelly-like paste was transferred onto the 3D-printed mould and pressed into pellets between the parafilm sheet and backing paper.

**Video 2. Free-feeding adult zebrafish consume drug pellets**.

Single-housed adult zebrafish were fed once daily with artemia during the day and given drug-pellets in the late afternoon. The video shows how zebrafish actively sought for and consumed the drug pellet voluntarily without any handling.

